# Improved estimation of hemodynamic response in fNIRS using protocol constraint and wavelet transform decomposition based adaptive algorithm

**DOI:** 10.1101/2020.04.25.062000

**Authors:** Sahar Jahani, Seyed Kamaledin Setarehdan

## Abstract

**Background:** Near infrared spectroscopy allows monitoring of oxy and deoxyhemoglobin concentration changes associated with hemodynamic response function (HRF). HRF is mainly affected by physiological interferences which occur in the superficial layers of the head. This makes HRF extracting a very challenging task. Recent studies have used an additional near channel which is sensitive to the systemic interferences of the superficial layers. This additional information can be used to remove the systemic interference from the HRF.

**New Method:** This paper presents a novel wavelet-based constrained adaptive procedure to define the proportion of the physiological interferences in the brain hemodynamic response. The proposed method decomposes the near channel signal into several wavelet transform (WT) scales and adaptively estimates proper weights of each scale to extract their share in the HRF. The estimation of the weights are done by applying data acquisition protocol as a coefficient on recursive least square (RLS), normalized least mean square (NLMS) and Kalman filter methods.

**Results:** Performance of the proposed algorithm is evaluated in terms of the mean square error (MSE) and Pearson’s correlation coefficient (R^2^) criteria between the estimated and the simulated HRF.

**Comparison with Existing Methods:** Results showed that using the proposed method is significantly superior to all past adaptive filters such as EMD/EEMD based RLS/NLMS on estimating HRF signals.

**Conclusions:** we recommend the use of WT based constraint Kalman filter in dual channel fNIRS studies with a defined protocol paradigm and using WT based Kalman filter in studies without any pre-defined protocol.

## I. INTRODUCTION

Neural activity of the brain results in releasing of vasoactive mediators leading to dilation of the surrounding arterioles and capillaries [1]. This increases the regional blood flow in the brain tissue which can be detected as an increase in the blood oxygenation level-dependent (BOLD) signal in the functional MRI (*f*MRI). This technique is used for brain function studies and analysis with many potential applications in clinics and researches. *f*MRI is, however, an expensive and non-portable equipment which makes it impossible to be used during normal daily activities. In comparison, functional near-infrared spectroscopy (*f*NIRS) which also monitors brain hemodynamics is a small, portable and a low-cost system [2][3]. *f*NIRS is based on the optical measurement technique that uses light in the near-infrared range (650-950 nm) to record variations in the brain blood flow during a mental activity. The light photons in the infrared range are either scattered by the soft tissue or absorbed by the oxy-Hb and deoxy-Hb within the blood. Recorded optical intensities are then used to calculate the changes in the concentration of these two main chromophores by the so-called modified Beer-Lambert Law (MBLL) [4]. Despite many advantages of fNIRS, the low amplitude hemodynamic response caused by mental activities is contaminated by the systemic or global physiological hemodynamic arising from heartbeat, breathing, and other homeostatic processes [5]. These natural hemodynamic occur in all body tissue including brain and superficial layers of the head. Several algorithms have been developed in the past to extract functional hemodynamic response from the systemic interferences. The most widely used and simplest method is low pass or band pass filtering (LPF or BPF)[6][7]. LPF and BPF are highly effective in removing cardiac oscillations ,but, since the frequency content of the hemodynamic response overlaps with the systemic interference, these methods are partially ineffective. Another common method is Block Averaging (BA) [8][9] with assumption that there is a phase difference between physiological components of each stimuli. In this technique a large number of trials of *f*NIRS recordings are needed to estimate hemodynamic response function (*HRF*) satisfactorily. Other methods for HRF extraction are based on principle component analysis (PCA) [6][10], independent component analysis (ICA) [11][12] and general linear model (GLM) [13]. In both ICA and PCA methods, the difference between spatial distribution of the HRF and the systemic interferences are considered. In PCA [14], the physiological interferences are reduced by removing the first principle component of N-channel signals. In ICA [15], the spatial distribution of independent components were used to identify the superficial components. Although both PCA and ICA techniques have improvement in signal gain, a main disadvantage of them is that the noise propagates from noisy channels to the other channels, resulting in underestimating the HRF. GLM has also shown to be beneficial in terms of signal gains, but it depends heavily on the selection of the basis functions. More recent promising method for reduction of physiological interference from fNIRS measurements is the use of the so-called “reference channel” [16][17][18][19][20][21][22][23]. The reference channel with the source-detector (SD) separation distance (<1) is an additional channel that is only sensitive to hemodynamic fluctuations within superficial layers. In [21] Saager et al., take the reference channel as systemic interference and cancel it from far distance source detector by linear minimum mean square (LMMSE). Other studies [16][22][23][24][25] have used adaptive filters to reduce systemic interferences. In [22][26] Zhang et. al employed the empirical mode decomposition (EMD) technique to decompose the near channel into its intrinsic mode functions (IMFs). Then, proper weights for each IMF were determined by means of an adaptive filter to estimate the weights of the systemic interference in the signal of the far channel (SD separation ≈ 3cm). In continue, the so-called problem of “mode mixing” was identified as the main drawback of the EMD method [27]. In an effort to reduce the effect of this drawback, the Ensembled EMD (EEMD) technique was developed by Wu et al. [28]. An adaptive algorithm based on the EEMD technique was applied to the dual channel *f*NIRS system in order to improve the cancelation of the systemic interference from HRF [23]. It has also been showed that EEMD based adaptive filters outperform HRF estimation in comparison with BA, BPF, PCA and ICA [23]. Static estimators such as recursive least square (RLS) and least mean square (LMS) are commonly used filters to estimate the weights of the systemic interference in the signal of the far channel [29][22][23][24]. In [17][30], the authors employed dynamic estimators such as state-space modeling with Kalman filter to estimate the *HRF*. Although, all these techniques improve the quality of the extracted HRFs from the raw signal of the far channel, but, we believe that there can be more improvement in the quality of the extracted signals if we include the priory information of the data collection paradigm in the extraction procedure. fNIRS studies for performing a cognitive task usually include a data acquisition protocol that contains a time sequence of applying the rest and task periods.

In this work, we applied the information of the stimulus times as a constraint on an adaptive filtering procedure to avoid unsuitable weights for the regressors. In addition, instead of EMD and EEMD, we employed the wavelet transform (WT) in order to decompose a signal into its orthogonal components [31]. This is a suitable property when we feed information into an adaptive filter. Hence, in this study, we propose two main ideas to improve the quality of the estimated HRFs: 1) decomposing the signal of the near channel into its constituents by WT, and 2) applying a protocol constraint to the RLS, LMS and Kalman filters for finding suitable weights for the decomposed components of near channel. To demonstrate the effectiveness of the proposed algorithm, we used a semi-real fNIRS data set which was produced by adding a synthetic HRF (true HRF) to real rest state data of far channel. The extracted HRFs by the proposed algorithm were then compared to the true HRF using the mean square error (MSE) and Pearson’s correlation (R^2^) parameters as quantitative criteria. In continue, the details of the proposed algorithm along with introducing the data set are explained in section II. Section III presents the results obtained by the proposed method for the data which was collected over 12 healthy subjects and compares tahem to those of the existing methods. Finally, Sections IV and V present the discussions and conclusions respectively.

## II. MATERIALS & METHODS

The schematic of the configuration of a dual channel fNIRS probe along with the block diagram of the proposed novel algorithm for HRF extraction is shown in Figure1. The light source (S) transmits photons into the head and the two detectors (D1 and D2) collect part of the scattered photons which travel the banana-shaped pathways through the cortex and scalp/brain tissues.

**Figure 1).**
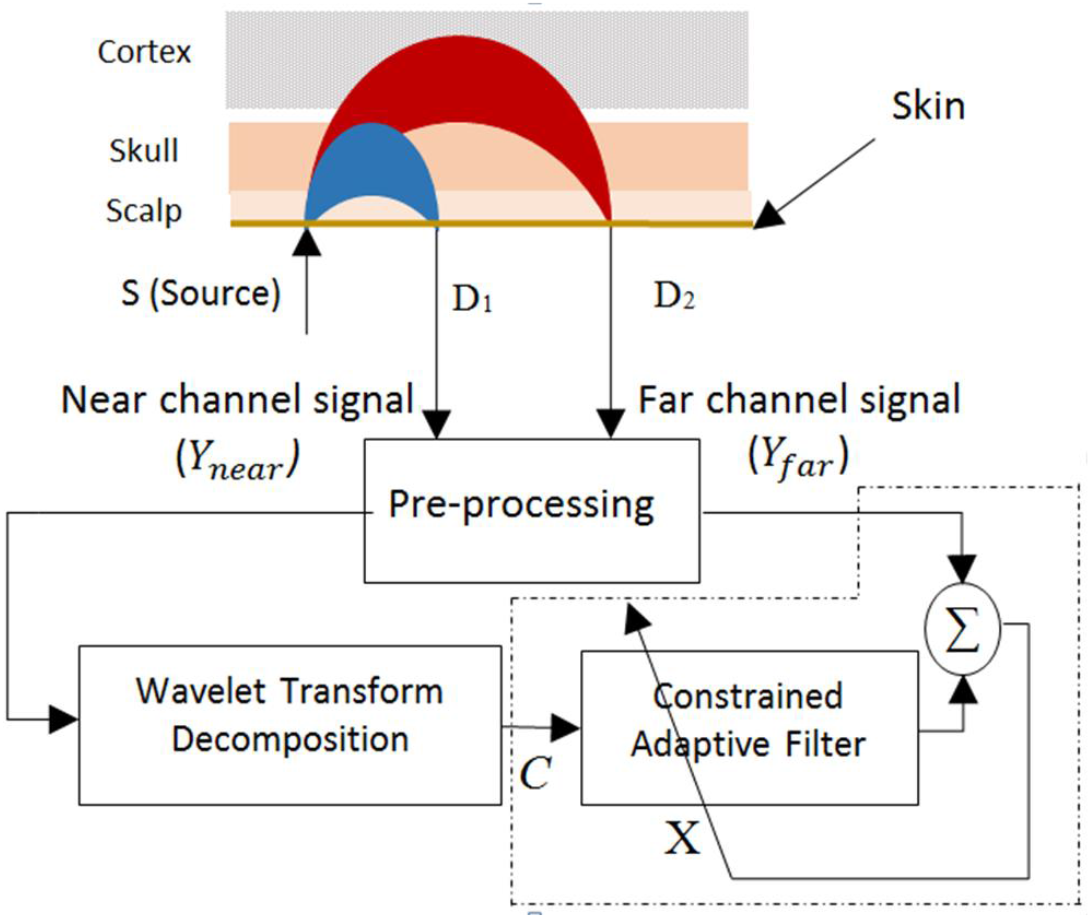
Schematic of the configuration of a dual channel fNIRS probe and its output to the block diagram of our proposed method.

Based on this configuration there are two channels of near (S-D_1_) and far (S-D_2_), the signals of which will go through signal pre-processing steps (section II.e). In this configuration, we set the inter SD distances for the near and far channels to <1cm and ≈3cm respectively. Previous studies have stated that the penetration depth of light photons in the brain tissue is about half of the SD distance [13]. Therefore, the near channel only contains information from the hemodynamic changes of the shallow tissue (skin and scalp) that includes heart rate, breathing, Mayer, and low-frequency oscillations, while the far channel includes hemodynamic of cortex which partially includes all these plus the hemodynamic of the functional activity of the brain. In our formulation we refer to the signal of near channel as *Y*_*near*_ and the signal of the far channel as *Y*_*far*_ with the following relationship:

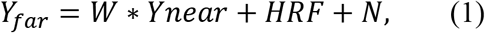

where *W* is the weight to be identified, HRF is the hemodynamic response function, and *N* is the measurement noise. By rearrangement equation (1) can be rewritten as:

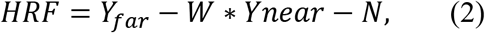

which emphasizes the fact that *HRF* can be derived by subtracting the estimated weight of physiological interference *Y*_*near*_ from the signal recorded at the far channel *Y*_*far*_.

In this work, instead of simply subtracting the weighted *Y*_*near*_ signal from the *Y*_*far*_ signal, we subtract the weighted wavelet transform scales of the *Y*_*near*_ signal from the *Y*_*far*_ signal [22] as shown in equation (3).

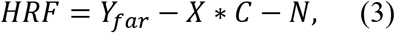

where *C* is the wavelet transform scales of the *Y*_*near*_ signal, *X* is the weight vector of the wavelet transform scales to be identified (to be adjusted) by the constrained adaptive filter as shown in Fig. 1. In continue the proposed novel algorithm is described in more details.

### a. Near channel signal decomposition

As it was mentioned before, instead of simply subtracting the weighted *Y*_*near*_ signal from the *Y*_*far*_ signal, we used the weighted wavelet transform scales of the *Y*_*near*_ signal for a more effective removal of the systemic interferences from the *Y*_*far*_ signal. In this study, we employed the orthogonal and compactly supported *Daubechies 7* mother wavelet [32] in order to decompose the HbO concentration signal of the near channel into its scales up to 10 levels (*d*_1__*d*_10_) using the wavelet package of Matlab software (The MathWorks Inc., Natik, MA, USA). An example of the decomposition of the HbO concentration signal of the near channel is shown in Figure 2.

**Figure 2).**
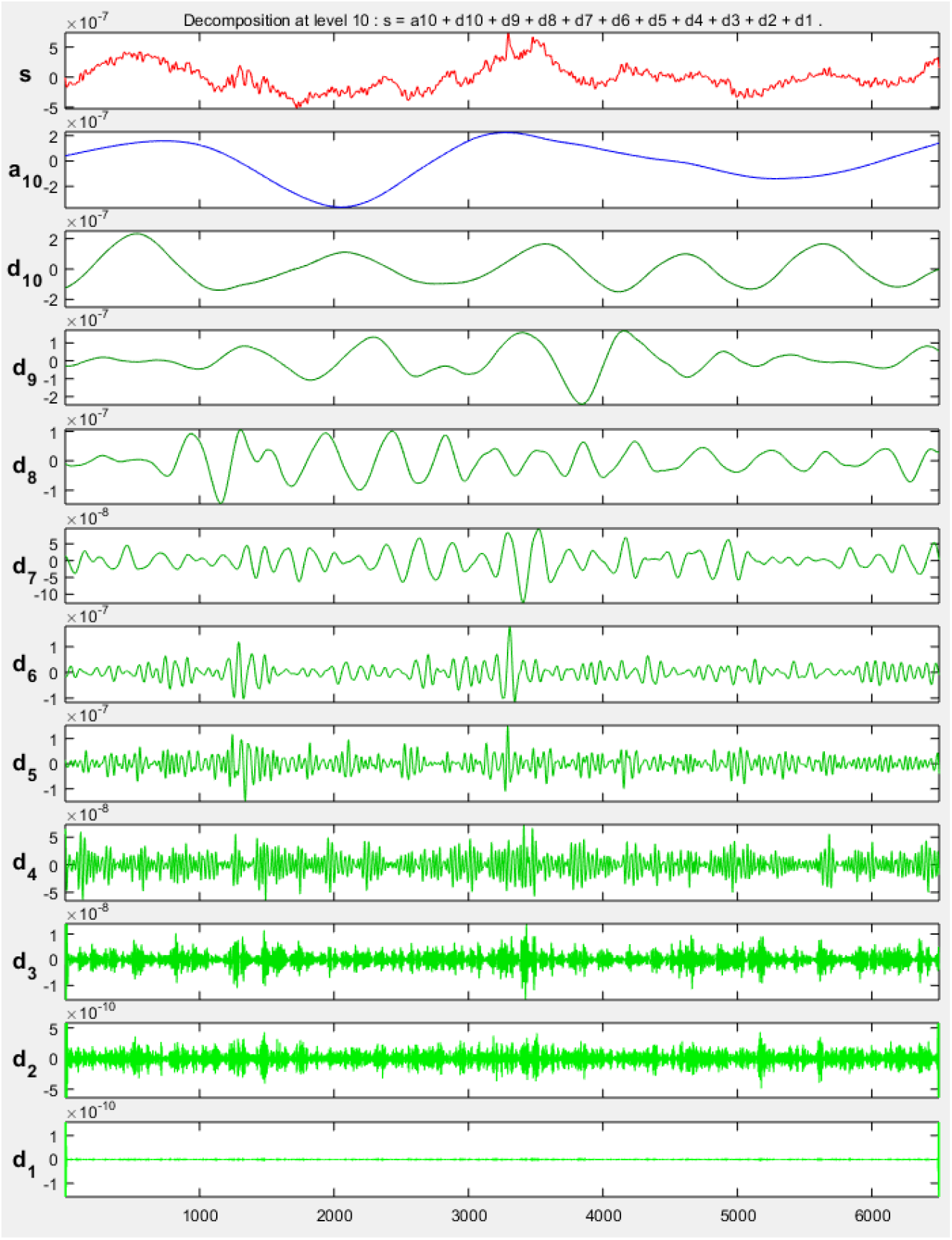
Example of the decomposition of the HbO signal of the near channel, The original signal (s), the approximation component (a10) and the details components (d_1__d_10_).

The effectiveness of the proposed wavelet transform based algorithm highly depends on the selection of the proper weights. For this reason, the weights vector X is estimated by the constrained adaptive filtering method as shown in Figure 3.

**Figure 3).**
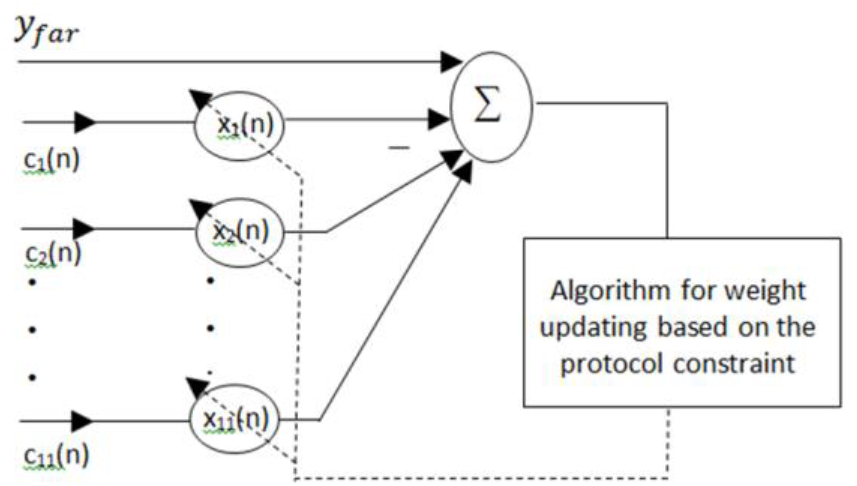
Constrained adaptive filtering method, the details of dashed box in Figure 1.

As seen in Figure 3, the near channel signal is decomposed into its scales by the wavelet transform producing L= 11 new signals of *c*_1_(*n*) = *a*_10_ (the approximation signal) and *c*_*i*_(*n*)|_*i*=2:11_ = *dj*|_*j*=*i*−1_ (the detail signals), where *n* is the signal samples (time index) and *i* is the scale index (*i* = 2 *to L*). Each *c*_*i*_ signal is then weighted by the corresponding weight of *x*_*i*_ and the 11 weighted signals are then summed and the result is subtracted from the corresponding far channels signal (equation 3) producing the HRF signal. In this study, three different techniques of *Normalized least mean squares* (NLMS), *Recursive Least Squares* (RLS) and *Kalman filter* are used in the weight estimation block, the results of which are then compared. In continue, each of these methods is explained in more details.

### b. Constrained Adaptive Filtering Techniques

#### NLMS filter

In Least Mean Squares (LMS) based adaptive filtering the filter weights are dynamically updated in order to minimize the mean squared error between the y_far_ signal and the sum of the weighted wavelet transform scales of the y_near_ (C). Since the LMS is highly sensitive to the amplitude of its input, we used the Normalized LMS (NLMS) instead, were the two signals are normalized with respect to their standard deviations as follows:

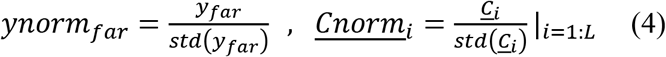

The error in the NLMS algorithm can be formulized as follows:

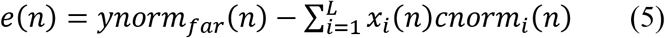

where *i* is the wavelet decomposition scale number and *n* is data samples. The filter coefficients *x*_*i*_(*n*) are updated via the steepest descent method as follows:

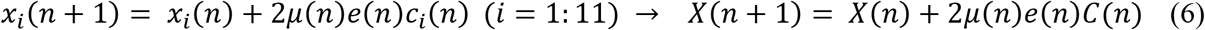

The parameter 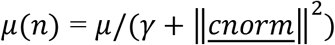 is the step size that controls the convergence rate of the adaptive filter. Parameter *γ* is a small positive number which is used to prevent dividing by zero. The initial values for *μ*(0) and <u>*X*</u>^*T*^(0) were set to 10^−4^ and [1 0 0 0 0 0 0 0 0 0 0] respectively.

#### RLS filter

The RLS algorithm exhibits extremely fast convergence while increases the computational complexity. RLS can recursively find the coefficients that minimize a cost function which is the sum of the weighted least squares as it is mentioned in follows:

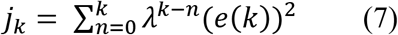

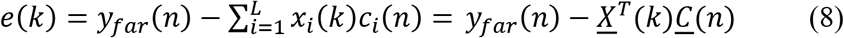

where *j*_*k*_ is cost function of RLS adaptive filter and λ is the forgetting factor.

To calculate the optimized weight vector that minimizes the cost function, we calculate derivate 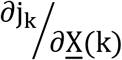, k = 0, … , n − 1, which is then set to zero. This produces the optimized weight vector as follow:

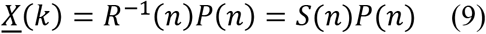

where *R*(*n*) is the autocorrelation of the wavelet decomposed signal of near channel and *P*(*n*) is the cross correlation between the *y*_*far*_ and the wavelet decomposed signal of near channel. Finally, the updating weight vector for wavelet decomposition signals is as follow:

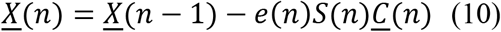

The initial value for *S*(*n*) was set to *S*(0) = *δ* × *I*_*L*×*L*_, where *I*_*L*×*L*_ is an identity matrix and *δ* is the regularization parameter. The weight values were initialized to <u>*X*</u>^*T*^(0) = [0 0 0 0 0 0 0 0 0 0 0].

#### Kalman filter

The Kalman filter is an optimal and dynamic estimator where the new measurements can be processed as they arrive. In the case of Gaussian noise, knowing the mean and standard deviation of the noise, the Kalman filter is the best linear estimator that minimizes the mean square error of the estimated parameters. A Kalman filter projects the measurements onto the state of a discrete-time process that is governed by the linear stochastic difference equation and with a measurement as:

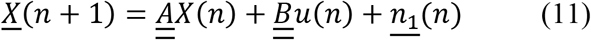

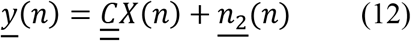

where matrix 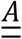 is the state at the previous time step (n) to the state at the current step (n+1) and matrix 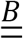 relates the optional control input to the state *X*. The matrix 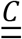 in the measurement equation relates the state to the measurement <u>*y*</u>(*n*).

The random variables <u>*n*_1_</u> (*n*) and <u>*n*_2_</u> (*n*) represent the process and measurement noise that are assumed to be white, independent from each other, and with normal probability distributions. The process noise covariance and measurement noise covariance matrices might change with each time step or measurement, however they are assumed constant:

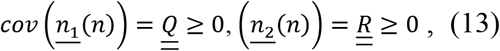

The process noise value determines the variation of the states at each time step. Considering a small value for process noise will approach the estimator to the static estimator, and a large one will significantly vary estimator over time. In this study, the process noise covariance diagonal terms were set to 10^−6^ and the diagonal terms of measurement noise were set to 10^−2^.

The equation that computes an *a posteriori* state estimate 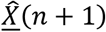 as a linear combination of an *a priori* estimate 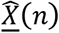 and measurement innovation that is a weighted difference between a measurement <u>*y*</u>(*n*) and the measurement prediction 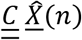 is as follows:

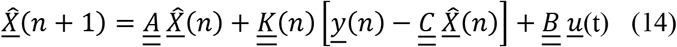

where 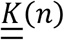 is the *blending factor* that minimizes the *a posteriori* error covariance equation. One form of 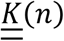 is:

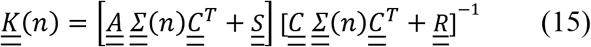

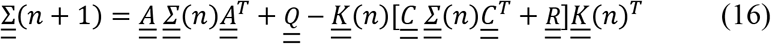

The Kalman filter initialization of the state vector estimate 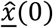 and estimated state covariance 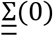 was set to:

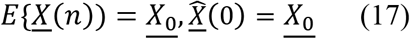

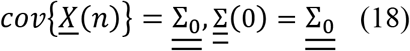

In this study, <u>*X*_0_</u> was set to the obtained values by RLS and 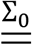 was set to a 10^−4^ × *I*, where *I* is the identity matrix.

### c. Constrained LMS, RLS and Kalman filter

Acquisition of *f*NIRS data is usually carried out based on and during a predefined protocol that consists of several rest and mental tasks time intervals. Estimation of functional hemodynamic response during mental task time intervals has been the aim of many previous studies [20][22][23]. In all these studies the influence of the rest and task time intervals on the updating coefficients of filters for regressing out the physiological noises are considered the same. As a matter of fact since there is no brain activity during the rest intervals, therefore, it is expected that the signals of the near and far channels contain only the physiological interferences. On the other hand, it is expected that during the mental task intervals, signals of the two channels are different, hence, the mutual information between near and far channels become more dissimilar. We propose to use the amount of the dissimilarity between the two channels as an information source to update the weights of the adaptive filters by considering an extra coefficient in the adaptive filter. This idea can be explained as follows. By the stimulus onset time, the corresponding hemodynamic response appears after a period of 8 to 10 seconds in the signal and by the end of the stimulus, the hemodynamic response disappears by a delay of 8 to 10 seconds. Since the adaptive filter tries to reduce the difference between the near and far channel signals, this extra coefficient will show a trend opposed to the hemodynamic response. When this coefficient is normalized between 0 and 1, its trend would be like the one which is depicted in Figure 4.

**Figure 4.**
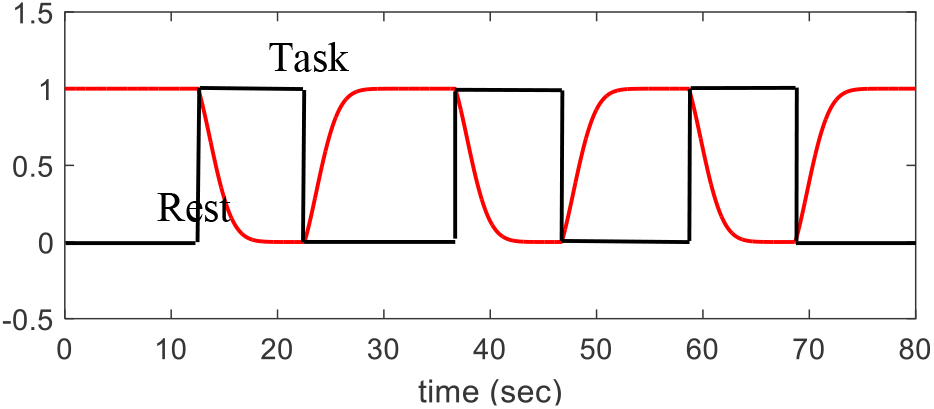
The rests and tasks blocks (black), the trend of protocol coefficient value (red).

By applying the protocol constraint *α*(*n*), the weight updating strategies for the constrained NLMS (CNLMS) Eq.19, constrained RLS (CRLS) Eq.20, and constrained Kalman (CKalman) Eq.21 filters should be modified as:

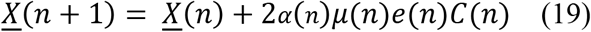

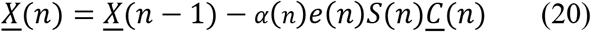

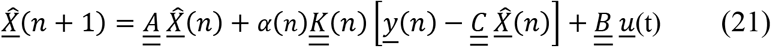

### d. Subjects and data acquisition

For performance analysis of the proposed algorithm 12 healthy subjects (age mean ± standard deviation: 26±8 years) were selected for data acquisition. Each subject completed a questionnaire to provide demographic information, drug use history, and physical status. A continuous wave *f*NIRS system with dual channel probe that is developed and evaluated by our team at the University of Tehran [33][34] is used. The probe, consists of 4 channels (2 near (SD<1cm) and 2 far (SD≈3cm) channels), was placed on the participants’ prefrontal cortex area. The raw optical intensity values at two different wavelengths (730nm and 850nm) were recorded by the fNIRS system with a sampling frequency of 50 Hz. During data acquisition procedure, the participants were asked to sit back relaxed in a dark silent room and have a 2 minutes rest.

### e. Preprocessing

In the preprocessing step, the intensity signals of the far and near channels are initially converted into the concentration signals of HbR and HbO by means of the Modified Beer-Lambert Law [4]. Next, the synthetic simulated HRF-HbO and HbR were respectively added to the HbO and HbR of far channel signals. The HRF inter stimulus intervals (the intervals with zero concentration value in between each two HRFs) were taken from a uniform distribution (rand) over the period of 5 to 10 seconds which provided 5 to 8 stimuli during ~2-min length of the data. An example of a synthetic true HRF is shown in Figure 5.

**Figure5.**
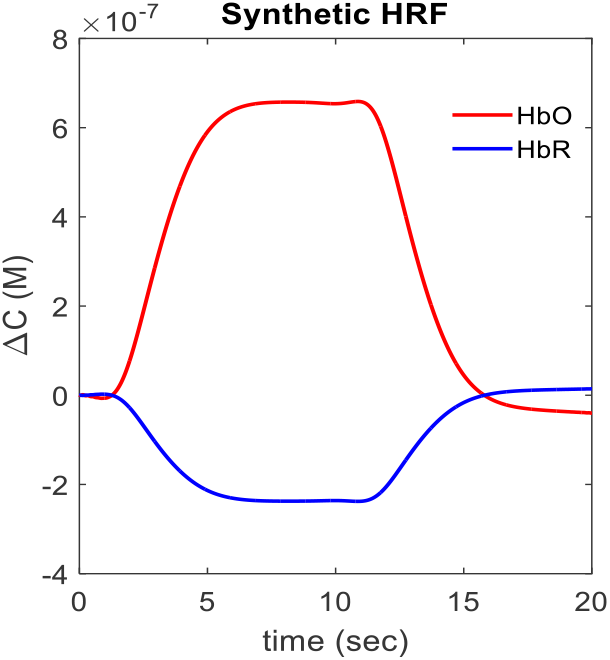
An example of a synthetic simulated HRF.

The concentration signals of HbO and HbR before adding synthetic HRF (during resting state) and after adding synthetic HRF for far and near channels are depicted in Figure 6 a-d.

**Figure 6.**
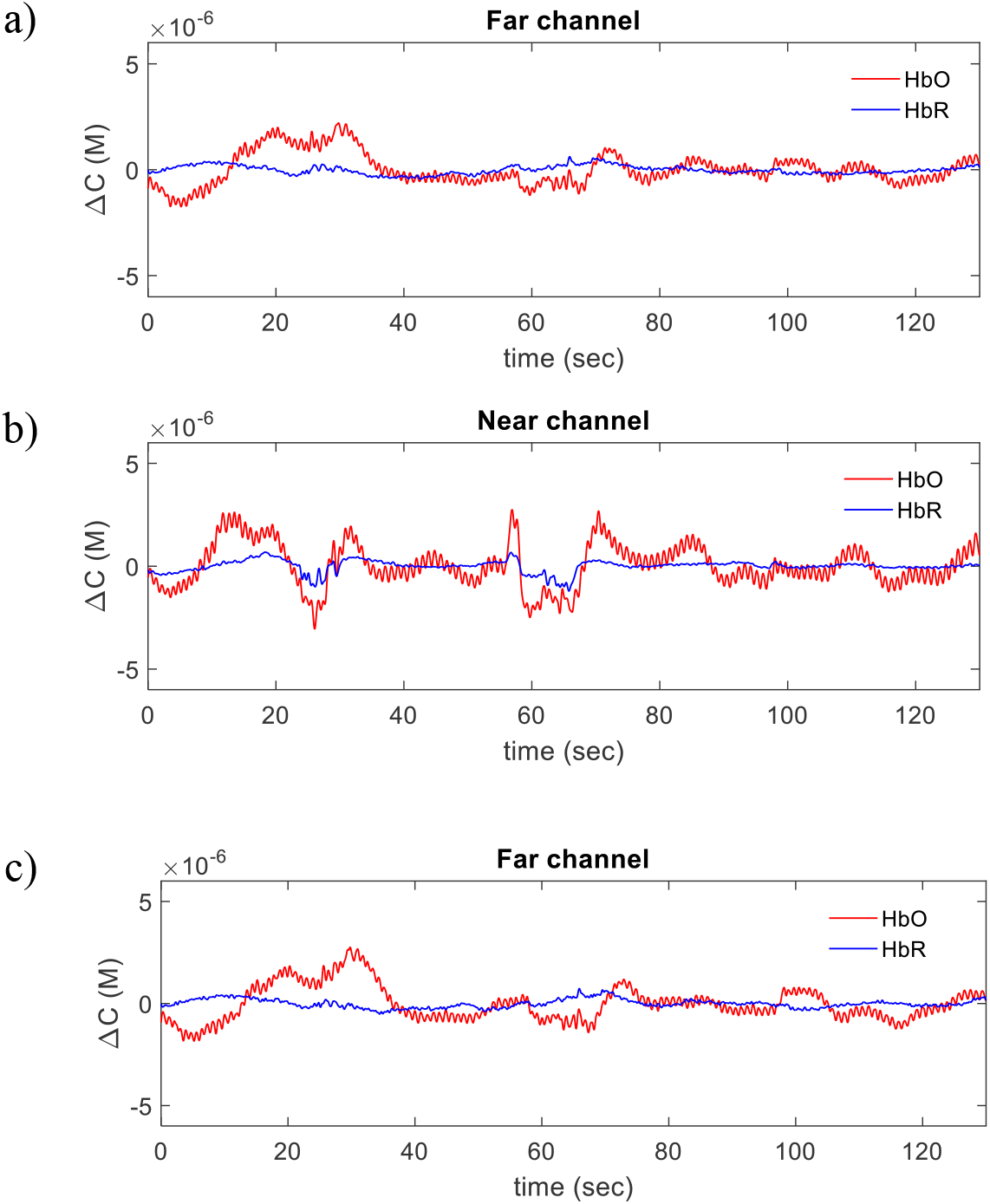
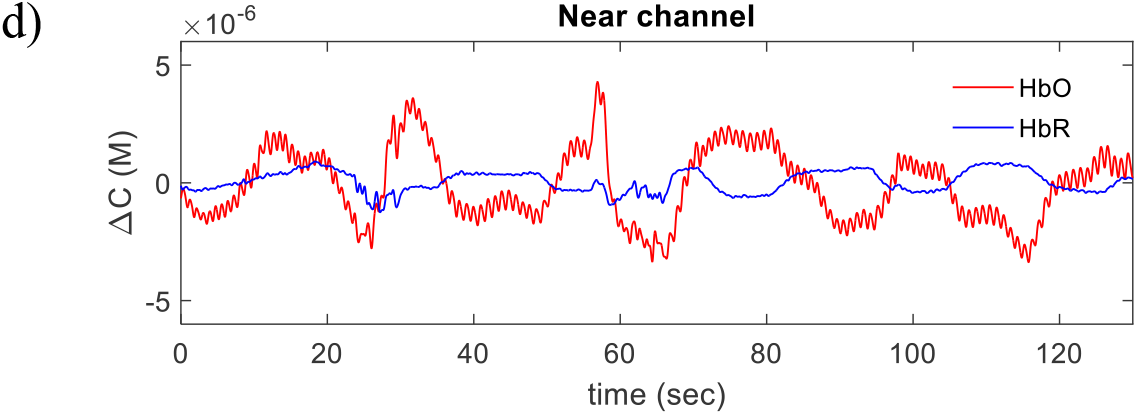
HbO and HbR concentration signals of real rest state signal before adding synthetic HRFs to a) far channel, b) near channel, after adding synthetic HRF to c) far channel and d)near channel.

Then, the concentration signals of HbO and HbR of the far and near channels are band pass filtered with a 3^rd^ order Butterworth filter (bandwidth 0.005-2 Hz) to remove both the trend of the signals and the high-frequency noises. These filtered signals are then used in the rest of the process as Y_near_ and Y_far_.

## III. RESULTS

### a. Validation and Comparing the proposed algorithm with other existing methods

To estimate the HRF signal by the proposed method and compare it with other previously mentioned algorithms, the Y_near_ signals were first decomposed into their components by three different methods of the wavelet transform, EMD and EEMD. The number of wavelet decomposition levels was set to 10 [31] and the number of decomposition levels for EMD and EEMD were set to 9 by their intrinsic criterion [22][23]. Then the decomposed data of Y_near_ were used for updating the weights of CNLMS, CRLS and CKalman filters. By extracting the appropriate weights for each decomposed signal of Y_near_, the estimated HRF can be easily measured by Eq.3. To smoothen and de-noising the results, the Savitzky-Golay filter with a polynomial order of 3 and frame size of 31 sample points is used [35]. Finally, the estimated HRFs, which are obtained by the WT, EMD and EEMD based NLMS, RLS and Kalman filter, are block averaged over the specific intervals of the trials. The true synthetic HRF and estimated HRF after block averaging are shown in Figure 7.

**Figure 7.**
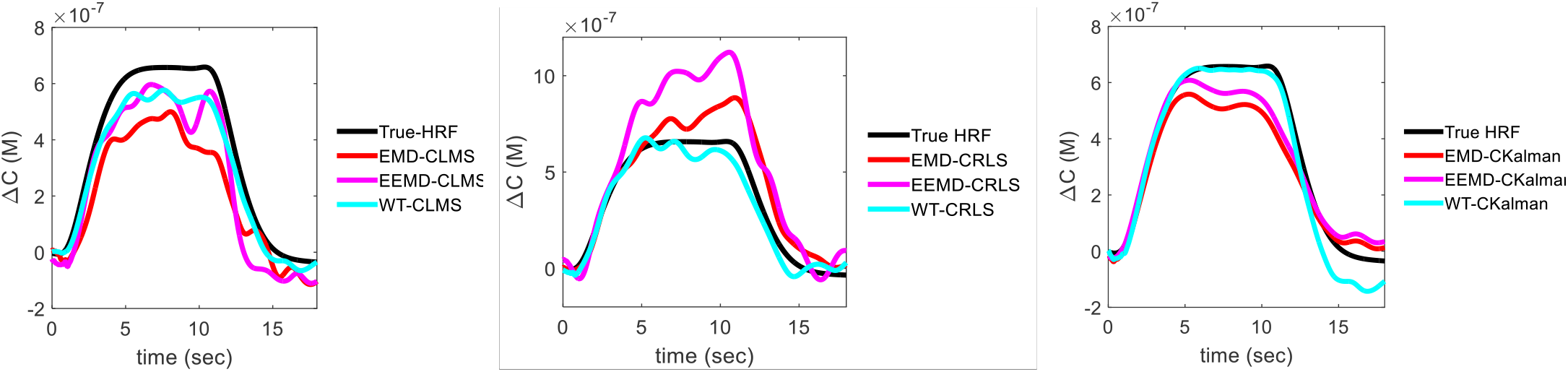
The block averaging of estimated HRFs by the mentioned algorithms

To compare the performance of the proposed method to those reported in the literature [23][22][16][36] the two quantitative criteria of: 1) mean squared error, 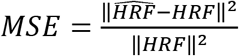, and 2) square of Pearson correlation coefficients (R^2^) are used in this study. R^2^ metric is obtained by Matlab software (Matlab function: corr, The MathWorks Inc., Natik, MA, USA).

The results of calculating MSE and R^2^ between the true and the estimated HRFs over all subjects are summarized in Table 1 and also are shown as bar graphs in

**Table 1.** Mean and standard values of MSE and R^2^ obtained by different HRF extraction algorithms.

| MSE | WT | EEMD | EMD |
| --- | --- | --- | --- |
| CKalman | 0.164±0.08 | 0.23±0.1 | 0.27±0.13 |
| Kalman | 0.19±0.1 | 0.27±0.11 | 0.29±0.14 |
| CRLS | 0.215±0.1 | 0.345±0.14 | 0.35±0.14 |
| RLS | 0.261±0.12 | 0.38±0.2 | 0.48±0.24 |
| CNLMS | 0.241±0.11 | 0.397±0.2 | 0.452±0.2 |
| NLMS | 0.273±0.12 | 0.434±0.2 | 0.45±0.2 |

| $R^2$ | WT | EEMD | EMD |
| --- | --- | --- | --- |
| CKalman | 0.93±0.03 | 0.9±0.05 | 0.9±0.04 |
| Kalman | 0.9±0.04 | 0.88±0.06 | 0.87±0.06 |
| CRLS | 0.91±0.04 | 0.84±0.05 | 0.82±0.1 |
| RLS | 0.88±0.05 | 0.84±0.09 | 0.8±0.12 |
| CNLMS | 0.87±0.06 | 0.82±0.09 | 0.81±0.1 |
| NLMS | 0.84±0.07 | 0.8±0.1 | 0.81±0.1 |

Figure 8.a and b. Moreover, the MSE and R^2^ between the mentioned adaptive filters with protocol constraint and without it are compared respectively. The bars represent the mean and the error bars symbolize the standard deviation. As it can be seen, the WT-CKalman method could outperform all the other methods in reducing the MSE and enhancing R^2^.

**Figure 8.**
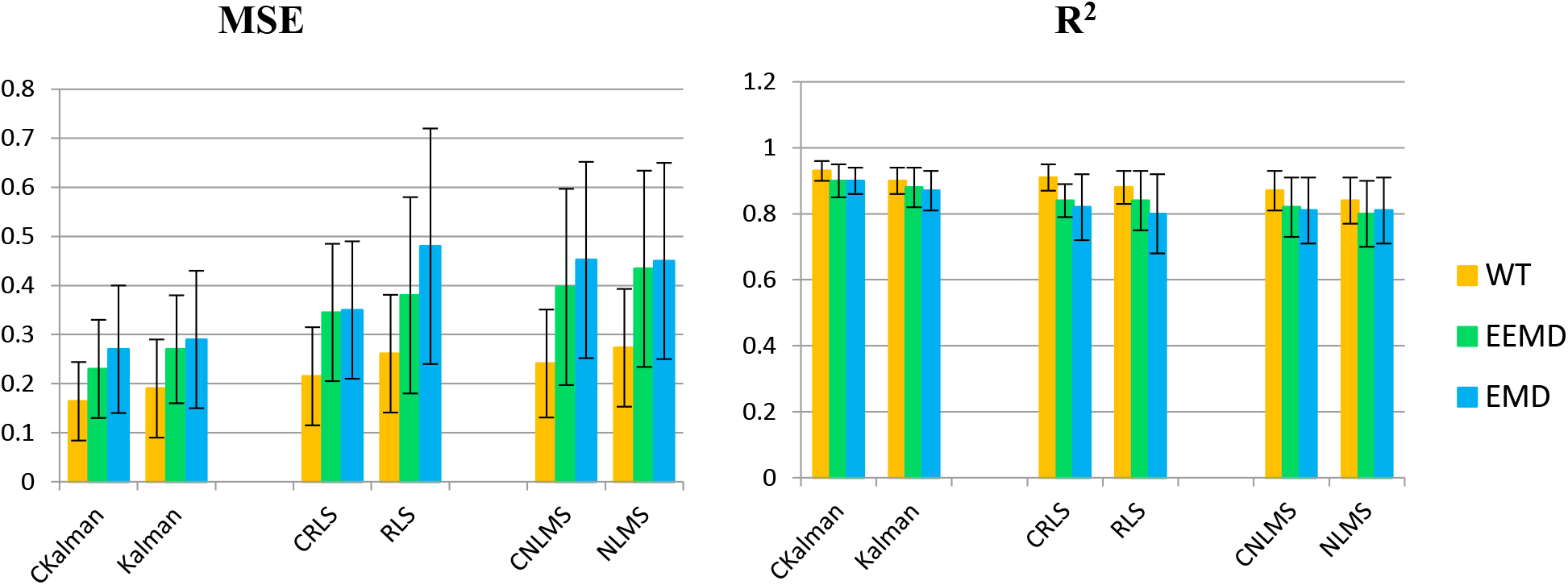
a) mean squared errors (MSE) b) pearson R^2^ coefficients, between simulated and true *HRF*. The bars represent the mean and the error bars symbolize the standard deviation over all subjects.

In continue, the Paired student’s t-tests were used to evaluate statistically significant differences in HRF extraction between different methods (criteria for significance: p-value<0.01 (**) and also p-value<0.05 (*)). The p-values for significant difference between MSE and R2 values of the aforementioned HRF extraction algorithms across all subjects are summarized in Tables 2 and 3. The MSE obtained for WT-CKalman was significantly lower than that of the other methods, except those obtained by WT-Kalman. Also, WT-CKalman showed significantly higher R2 values than all the other mentioned algorithms except than WT-CRLS. Also, it can be observed that using the WT for signal decomposition instead of the EEMD and EMD methods shows significant improvement in MSE for all kind of adaptive filters. This also resulted in a significant enhancement in R^2^ for nearly most of algorithms except for the WT_RLS and WT-Kalman compared to EMD-RLS and EEMD-RLS algorithms respectively. The other novel approach besides using WT at the decomposition level is using the information of data acquisition paradigm as a constraint on updating weights of the adaptive filters. With regards to the calculated p-values in Tables 2 and 3, it can be concluded that using the paradigm information could resulted in a better MSE and R^2^ value for most aforementioned algorithms.

**Table 2.** The p-values for significant difference between MSE values of different HRF extraction algorithms. p<0.01 (**),p<0.05 (*), not significant (N)

| MSE | WT-CKalman | EEMD-CKalman | EMD-CKalman | WT-Kalman | EEMD-Kalman | EMD-Kalman | WT-CRLS | EEMD- CRLS | EMD- CRLS | WT-RLS | EEMD- RLS | EMD- RLS | WT- CNLMS | EEMD- CNLMS | EMD- CNLMS | WT- NLMS | EEMD- NLMS |
| --- | --- | --- | --- | --- | --- | --- | --- | --- | --- | --- | --- | --- | --- | --- | --- | --- | --- |
| WT-CKalman |  |  |  |  |  |  |  |  |  |  |  |  |  |  |  |  |  |
| EEMD-CKalman | * |  |  |  |  |  |  |  |  |  |  |  |  |  |  |  |  |
| EMD-CKalman | ** | N |  |  |  |  |  |  |  |  |  |  |  |  |  |  |  |
| WT-Kalman | N | * | ** |  |  |  |  |  |  |  |  |  |  |  |  |  |  |
| EEMD-Kalman | ** | * | N | ** |  |  |  |  |  |  |  |  |  |  |  |  |  |
| EMD-Kalman | ** | * | * | ** | N |  |  |  |  |  |  |  |  |  |  |  |  |
| WT-CRLS | * | N | ** | N | * | ** |  |  |  |  |  |  |  |  |  |  |  |
| EEMD- CRLS | ** | * | ** | ** | ** | * | ** |  |  |  |  |  |  |  |  |  |  |
| EMD- CRLS | ** | * | ** | ** | ** | * | ** | N |  |  |  |  |  |  |  |  |  |
| WT-RLS | ** | * | N | * | N | N | * | ** | ** |  |  |  |  |  |  |  |  |
| EEMD- RLS | ** | * | ** | ** | ** | ** | ** | N | N | ** |  |  |  |  |  |  |  |
| EMD- RLS | ** | * | ** | ** | ** | ** | ** | ** | ** | ** | ** |  |  |  |  |  |  |
| WT- CNLMS | ** | N | * | * | * | * | N | ** | ** | N | ** | ** |  |  |  |  |  |
| EEMD- CNLMS | ** | * | ** | ** | ** | ** | ** | * | * | ** | N | ** | ** |  |  |  |  |
| EMD- CNLMS | ** | * | ** | ** | ** | ** | ** | ** | ** | ** | ** | N | ** | * |  |  |  |
| WT- NLMS | ** | N | N | ** | N | N | * | ** | ** | N | ** | ** | * | ** | ** |  |  |
| EEMD- NLMS | ** | * | ** | ** | ** | ** | ** | ** | ** | ** | ** | * | ** | * | ** | ** |  |
| EMD- NLMS | ** | * | ** | ** | ** | ** | ** | ** | ** | ** | ** | N | ** | * | N | ** | N |

**Table 3.** The p-values for significant difference between R^2^ values of different HRF extraction algorithms. p<0.01 (**),p<0.05 (*), not significant (N)

| $R^2$ | WT-CKalman | EEMD-CKalman | EMD-CKalman | WT-Kalman | EEMD-Kalman | EMD-Kalman | WT-CRLS | EEMD-CRLS | EMD-CRLS | WT-RLS | EEMD-RLS | EMD-RLS | WT-CNLS | EEMD-CNLS | EMD-CNLS | WT-NLMS | EEMD-NLMS |
| --- | --- | --- | --- | --- | --- | --- | --- | --- | --- | --- | --- | --- | --- | --- | --- | --- | --- |
| WT-CKalman |  |  |  |  |  |  |  |  |  |  |  |  |  |  |  |  |  |
| EEMD-CKalman | * |  |  |  |  |  |  |  |  |  |  |  |  |  |  |  |  |
| EMD-CKalman | * | N |  |  |  |  |  |  |  |  |  |  |  |  |  |  |  |
| WT-Kalman | * | N | N |  |  |  |  |  |  |  |  |  |  |  |  |  |  |
| EEMD-Kalman | ** | N | N | N |  |  |  |  |  |  |  |  |  |  |  |  |  |
| EMD-Kalman | ** | * | * | * | N |  |  |  |  |  |  |  |  |  |  |  |  |
| WT-CRLS | N | N | N | N | * | * |  |  |  |  |  |  |  |  |  |  |  |
| EEMD-CRLS | ** | ** | ** | ** | * | * | ** |  |  |  |  |  |  |  |  |  |  |
| EMD-CRLS | ** | ** | ** | ** | ** | ** | ** | N |  |  |  |  |  |  |  |  |  |
| WT-RLS | ** | * | N | N | N | N | * | * | ** |  |  |  |  |  |  |  |  |
| EEMD-RLS | ** | ** | ** | ** | * | * | ** | N | N | * |  |  |  |  |  |  |  |
| EMD-RLS | ** | ** | ** | ** | ** | ** | ** | * | N | N | * |  |  |  |  |  |  |
| WT-CNLS | ** | * | * | * | N | N | * | * | ** | N | * | ** |  |  |  |  |  |
| EEMD-CNLS | ** | ** | ** | ** | ** | ** | ** | ** | N | ** | N | ** | ** |  |  |  |  |
| EMD-CNLS | ** | ** | ** | ** | ** | ** | ** | ** | N | ** | * | ** | ** | N |  |  |  |
| WT-NLMS | ** | ** | ** | ** | * | * | ** | N | N | * | N | * | * | N | * |  |  |
| EEMD-NLMS | ** | ** | ** | ** | ** | ** | ** | * | N | ** | * | N | ** | N | N | * |  |
| EMD-NLMS | ** | ** | ** | ** | ** | ** | ** | * | N | ** | * | N | ** | N | N | * | N |

## IV. DISCUSSION

*f*NIRS can be effectively employed to provide useful information for the study of cerebral activity. Since, the HRF of cerebral activity is highly degraded by physiological interferences; estimation of HRF is a challenging problem due to its small amplitude compared to the physiological components. Hence, in the present study, an effective method to improve the recovery of HRF from *f*NIRS signals is presented. In summary, we distinct HRF from physiological interference by means of dual channel *f*NIRS. First, the preprocessed signal of the near channel is decomposed to its different scales by the Wavelet Transform (WT); then, the weight of each component with distinct frequency content is adjusted by constrained adaptive filters to extract an estimate for physiological interferences. To evaluate and compare the proposed method with previous existing methods, synthetic HRF was added to real oxyhemoglobin changes of far channel, then the estimated HRFs were compared with the true simulated HRFs with two criteria of MSE and R^2^. As it is shown in Table 1, the CKalman method could outperform the other estimation methods in the sense of producing less MSE values. We believe this is the consequence of involving WT along with the protocol constraint into the estimation procedure which avoids the algorithm from falling into local optimums and extracts the estimated HRF quite similar to its simulated counterpart. The proposed method also produces the largest correlation between the recovered and simulated hemodynamic signals. This correlation delineates how well our method estimates the simulated signal details. In previous study, EEMD_RLS was introduced as the best among commonly used algorithm of BA, BPF, PCA and ICA [23]. Remarkably, the UB-CKalman method obtained the notable performance for all indexes with a significant difference compared to EEMD_RLS method. Due to the resulted p-values in Tables 2 and 3, WT-CKalman showed significantly lower MSE values than the other methods, except than of the WT-Kalman. It shows that combination of wavelet and Kalman can result in significant improvement for the studies where no protocol constraint is designated. Also, WT-CKalman significantly outperformed the other methods except than WT-CRLS in the sense of higher R^2^ values. This can be due to the use of protocol constraint idea since the WT_CRLS method has also resulted in significant improvement in R^2^ compared to the WT_RLS method. With regards to the resulting p-values in Tables 2 and 3, it can be concluded that the effect of using WT for signal decomposition and applying the information of data acquisition paradigm as a constraint on updating weights of the adaptive filters resulted in a better enhancement of both MSE and R^2^ in the most aforementioned algorithms.

We believe that the reason that makes the wavelet transform stand apart from the other transforms is its multi resolution nature. The constituents which make up the near channel signal ranges over multiple resolutions due to their sources. These components range from B-Waves, M-Waves and Respiration which belong to low frequency section, up to Heart beat which is treated as the high frequency component. Because Wavelet transform considers resolution of the signals in its decomposition procedure, it is more likely to decompose the near channel signal into components which are close to these real world signals.

Our method also produces lower variance in multiple runs of the algorithm which makes its results more reliable than other methods. We believe that this desired property arises from the fact that we have employed the transform and estimation methods which makes it more probable to extract signals with meaningful real world counterparts. As the hemodynamic signals are non-stationary [37], we also used an adaptive dynamic procedure in a hope to capture the non-stationary characteristics of the physiological components of the *f*NIRS signals. This makes our algorithm capable of estimating the time-varying weights for wavelet components which can be thought of time-varying parameters of the underlying model for the signals. This fact in conjunction with the fact that we impose no assumption on the amplitude, shape, and duration of the HRF signals make the whole algorithm robust against multiple sources of variations.

## V. CONCLUSION

In this study, we proposed a wavelet-based constrained adaptive dynamic filter (WT-CKalman) to regress out the physiological interferences from the brain hemodynamic response by using additional near source-detector information. The results of this study suggest that using wavelet transform instead of using EMD and EEMD outperforms the results of adaptive filters. Also, applying protocol constraint on dynamic (Kalman) and static (RLS and NLMS) adaptive filters resulted in better performance in reducing MSE and increasing R^2^ compared to the previous aforementioned methods. As a conclusion of this study, we recommend the use of WT-CKalman in dual channel fNIRS studies with a defined protocol paradigm and using WT-Kalman in studies without any pre-defined protocol.

## ACKNOWLEDGMENT

The authors would like to thank the Cognitive Sciences and Technologies Council (CSTC) of Islamic Republic of Iran for its support of this research work (contract No: 383, Date: 07/24/2016).

